# A role for Flipons and miRNAs in Promoter Specification during Development

**DOI:** 10.1101/2022.06.06.495008

**Authors:** Alan Herbert, Fedor Pavlov, Dmitrii Konovalov, Maria Poptsova

## Abstract

The classical view of gene regulation is based on prokaryotic models and the operon concept with protein-based transcription factors controlling the expression of metabolic pathways essential for bacterial adaptations in response to environmental changes. A new view for establishing cell identity is emerging in eukaryotes where RNA-based pathways provide the framework for the readout of genomic information. Another perspective poses that alternative DNA structures encoded by flipons enable switching of cellular responses from one state to another. Here we provide evidence that these RNA and DNA mechanisms are deeply connected. We present data supporting a model where flipons open up binding sites for microRNAs (miRNAs), leading to the establishment of bivalent promoters early in development whose location structures lineage-specific events. These outcomes are potentially influenced by ovarian and spermatozoan miRNAs, transmissions with evident evolutionary ramifications. The data supports a new perspective on genetic regulation, one in which the genome provides a canvas framed by flipons, sketched with miRNAs and embellished by proteins.

## Introduction

The approach we take follows that of Britten and Davidson who first proposed the programming of genomes by *trans*-RNAs that act at locations distant from where they are produced [1]. While there are many classes of trans-RNAs of varying sizes [2], we will focus on microRNAs (miRNAs) and examine the DNA sites to which they bind and their potential role in embryonic development. We posit that flipons, sequences that adopt non-B-DNA conformations under physiological conditions[3], play a role in targeting maternal and paternal miRNAs to particular locations to establish developmentally important bivalent promoters (BiPs), where both active histone 3, lysine 4 trimethylation (H3K4me3) and suppressive histone 3, lysine 27 trimethylation (H3K27me3) are present. The loss of one or other of these modifications at a later stage guides tissue formation and differentiation. We find that different flipon classes and evolutionarily conserved miRNA families seed sequences [4] colocalize with each other in promoter regions, along with the AGO1 and AGO2 argonaute proteins that are miRNAs effectors (AERs). The promoter so characterized are enriched for transcription factor genes that specify cell-fate.

### The Links between miRNAs and Gene Regulation

The ability of miRNAs to regulate phenotype was first shown by their effects due to their suppression of transcript translation. By changing the time window in which a protein was produced, such miRNAs alter the size and cellular composition of an organ formed during development [5]. In mice, knockout of conserved miRNAs has profound effects on embryonic development [4]. These miRNAs arise through a number of canonical and non-canonical processing pathways that output AERs [4]. In line with a role for *trans*-RNAs in these early outcomes, *Ago2* knockout is embryonically lethal with a failure to develop extra-embryonic tissue (Table 1). However, survival does not depend on the AGO2 enzymatic slicer activity needed to suppress translation as mice homozygous for a catalytically dead *Ago2*^*ADH*^ allele develop normally [6], suggesting that AGO2 is active through a different mechanism. Besides the cytoplasm, miRNAs also localize to the nucleus [2,7]. They modulate gene expression [2,8,9] and potentially a subset of alternative splices [10-12].

**Table 1.** Gene Ontogeny Enrichment for Promoters that have SIDDs paired with a Z-, G- or T-flipon 500 basepairs either side. Conserved miRNA seed-sequence location was not used in this analysis. (ER: enrichment ratio; TF: transcription Factor; Pol: Polymerase) (See Supplementary Data for additional details)

| Molecular Function |  |  |  |  |  | Biological Process (non-redundant) |  |  |  |  |
| --- | --- | --- | --- | --- | --- | --- | --- | --- | --- | --- |
| Flipon | Geneset | Description | ER | P-value | FDR | Geneset | Description | ER | P-value | FDR |
| Z-DNA | GO:0043565 | sequence-specific DNA binding | 4.65 | 0 | 0 | GO:0045165 | cell fate commitment | 7.70 | 1.78E-15 | 1.44E-12 |
|  | GO:0003700 | DNA-binding TF activity | 4.59 | 0 | 0 | GO:0001501 | skeletal system development | 4.48 | 2.32E-11 | 9.44E-09 |
|  | GO:0001067 | regulatory region nucleic acid binding | 4.86 | 0 | 0 | GO:0061564 | axon development | 4.46 | 1.44E-10 | 3.89E-08 |
|  | GO:0044212 | transcription regulatory region DNA binding | 4.87 | 0 | 0 | GO:0007389 | pattern specification process | 4.44 | 3.64E-10 | 7.38E-08 |
|  | GO:0000981 | DNA-binding TF, RNA Pol II-specific | 5.32 | 0 | 0 | GO:0048638 | regulation. Development and growth | 4.79 | 1.14E-09 | 1.85E-07 |
| G4 | GO:0043565 | sequence-specific DNA binding | 4.62 | 0 | 0 | GO:0045165 | cell fate commitment | 6.86 | 3.09E-13 | 2.51E-10 |
|  | GO:0003700 | DNA-binding transcription factor activity | 4.57 | 0 | 0 | GO:0050808 | synapse organization | 4.43 | 4.01E-10 | 1.63E-07 |
|  | GO:0001067 | regulatory region nucleic acid binding | 4.84 | 0 | 0 | GO:0007389 | pattern specification process | 4.30 | 7.46E-10 | 2.02E-07 |
|  | GO:0044212 | transcription regulatory region DNA binding | 4.86 | 0 | 0 | GO:0061564 | axon development | 4.15 | 1.55E-09 | 3.15E-07 |
|  | GO:0000976 | regulatory region DNA-specific binding | 5.00 | 0 | 0 | GO:0048568 | embryonic organ development | 4.03 | 2.88E-09 | 4.67E-07 |
| H-DNA | GO:0043565 | sequence-specific DNA binding | 5.20 | 8.27E-14 | 8.09E-11 | GO:0045165 | cell fate commitment | 9.19 | 1.54E-11 | 1.25E-08 |
|  | GO:0000981 | DNA-binding TF activity, RNA Pol II-specific | 6.31 | 8.80E-14 | 8.09E-11 | GO:0050808 | synapse organization | 5.46 | 3.00E-08 | 1.22E-05 |
|  | GO:0003700 | DNA-binding transcription factor activity | 5.21 | 7.07E-13 | 4.33E-10 | GO:0048568 | embryonic organ development | 4.65 | 6.65E-07 | 1.80E-04 |
|  | GO:0044212 | transcription regulatory region DNA binding | 5.37 | 3.16E-12 | 1.25E-09 | GO:0050890 | cognition | 5.69 | 1.23E-06 | 2.49E-04 |
|  | GO:0001067 | regulatory region nucleic acid binding | 5.35 | 3.40E-12 | 1.25E-09 | GO:0050803 | regulation. synapse structure/ activity | 6.10 | 1.78E-06 | 2.90E-04 |

### Flipons and Nucleosome Phasing

A role for Z-DNA and G4 flipons in gene expression has been proposed almost since the discovery of these non-B-DNA structures [13]. The flip to an alternative conformation requires energy, such as that liberated when polymerases and helicases hydrolyze ATP to separate DNA strands or that energy released instantaneously when nucleosomes are ejected from DNA (Figure 1A and B)[14]. The unwinding of the B-DNA helix due to these events generates negative supercoiling that drives the formation of left-handed DNA [15] and four-stranded quadruplexes (consensus sequence d(G_3_X_1-7_)_4_ [16-18]. The flip occurs faster than topoisomerases can relax the DNA torque [19]. The enrichment of Z- and G-flipons and the structures they form in promoter regions likely flags these sites for the cellular machinery to find easily, while also keeping transcription start sites (TSS) nucleosome-free [14,20,21].

**Figure 1.**
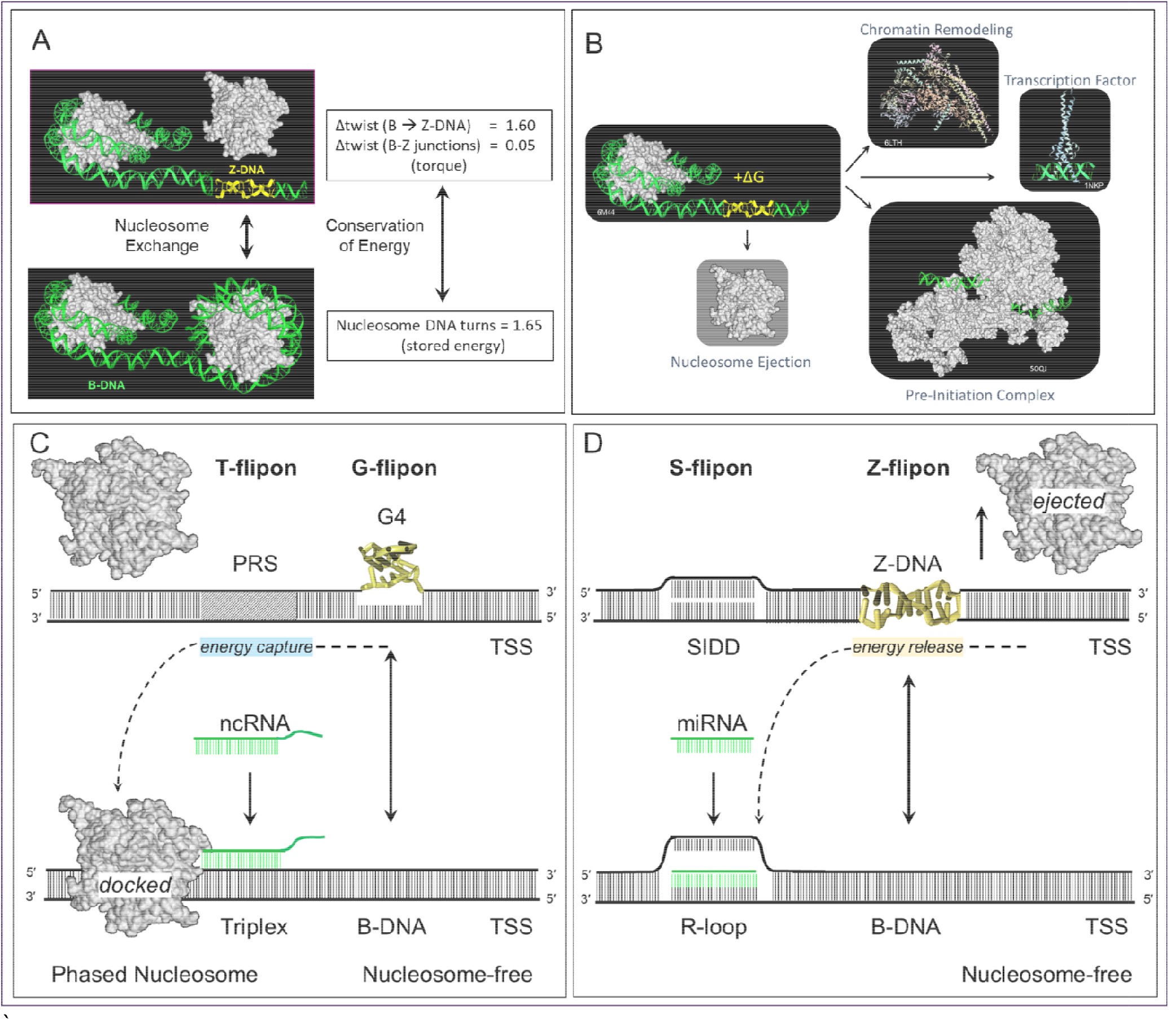
Converting torque into action **A**. The stress produced in underwound DNA is relieved by a flipon adopting the left-handed Z-DNA conformation or by wrapping the DNA around a nucleosome. The change in DNA twist when Z-DNA flips to B-DNA is equivalent to the number of turns in the solenoid formed by coiling the DNA around the histone core. The formation of Z-DNA can phase the placement of nucleosomes on DNA so that transcription start sites are accessible [54]. **B**. Release of the energy stored in nucleosome cores can power formation of different chromatin complexes that regulate chromatin state and the readout of information from the genome. **C**. Triplex formation by Purine Rich Sequences (PRS) within a T-flipon can also incorporate miRNAs and long non-coding RNAs [26] G-flipons are more stable than Z-DNA and require resolution by a helicase to generate unwound DNA that then wraps around nucleosomes to capture the free energy produced [20,21].**D**. The energy released by ejection of a nucleosome (indicated by upwards arrow) is captured by Z-DNA formation and accumulates until sufficient to power strand separation at the Stress Induced Sequence Destabilized (SIDD) sites that define S-flipons, allowing base-specific binding of ligands such as RNA, DNA and protein. In both cases, flipons are associated with nucleosome free regions and flag these sites for chromatin and transcriptional factors to bind, reducing the genomic search space.

These and other types of flipon can be detected *in cellulo* by using potassium permanganate (KMnO_4_) footprinting [22]. In particular, they identify sites of Stress Induced Duplex Destabilization (SIDDs) [23] that potentially bind *trans*-RNA to form R-loops [24,25] and triplexes [26,27]. Both of these alternative structures are incompatible with nucleosome assembly [28] or engagement of B-DNA specific proteins. SIDDs are AT-rich and are referred to here as S-flipons. Besides RNA and DNA, SIDDs can bind single-stranded DNA with high affinity, such as those KH domain proteins that regulate c-MYC gene expression [29] and alternative RNA splicing [30]. KMnO_4_ footprinting can also detect H-DNA, which is formed by the fold-back of a single-stranded region of polypurine or polypyrimidine DNA onto an adjacent sequence-matched duplex [22]. H-DNA represents a class of T-flipon that can also form triplexes with either *trans*-RNAs and DNAs [13,14].

The question arises whether the effects of *trans*-RNAs and flipons on gene expression are concerted. Here we present evidence in support of this possibility and for their joint role in early development.

## Results

We queried the flipon sequences identified by Kouzine et al. [22] in activated murine B cells by KMnO4 footprinting to assess their match to those seed sequences found in the set of miRNA families highly conserved in placental animals [4] (Figure 2A, Supplementary Data 1) (Z-DNA (n=25,059), G4 (n=20,253), SIDD (n=15,296), H-DNA (n=17,100)). We observed a frequent overlap of miRNA seed sequences with SIDDs (mean length =170 bps), in contrast to other flipon classes (Figure 2A). The most frequent overlap was with miR-203a (∼20%) that promotes epidermal differentiation by repressing the proliferative potential of dermal stem cells in mice, starting at embryonic day 13.5 (E13,5) [31]. Other miRNA matches were also common, including the miR-374 (4.56%) that is encoded within the X-inactivation center [32]. The high match frequency observed is consistent with a biological role for such interactions as the typical miRNA has 40-1500 copies per cell [33], a stoichiometry sufficient to cover many of the available 15,296 SIDD sites identified in the Kouzine data assuming that about half the miRNAs are nuclear [7]. There were a few instances of overlap of miRNA seed sequences with other flipons classes (Figure 2A). The Z-prone d(AC)_n_ sequence implicated in splicing [11,34] has a 4.5% overlap with miR-329/362. Other miRNA seed sequences containing d(G)_3_ (or its complement d(C)_3_) overlapped the annotation for G4-quadruplex (d(G_3_X_1-7_)_4_).

**Figure 2.**
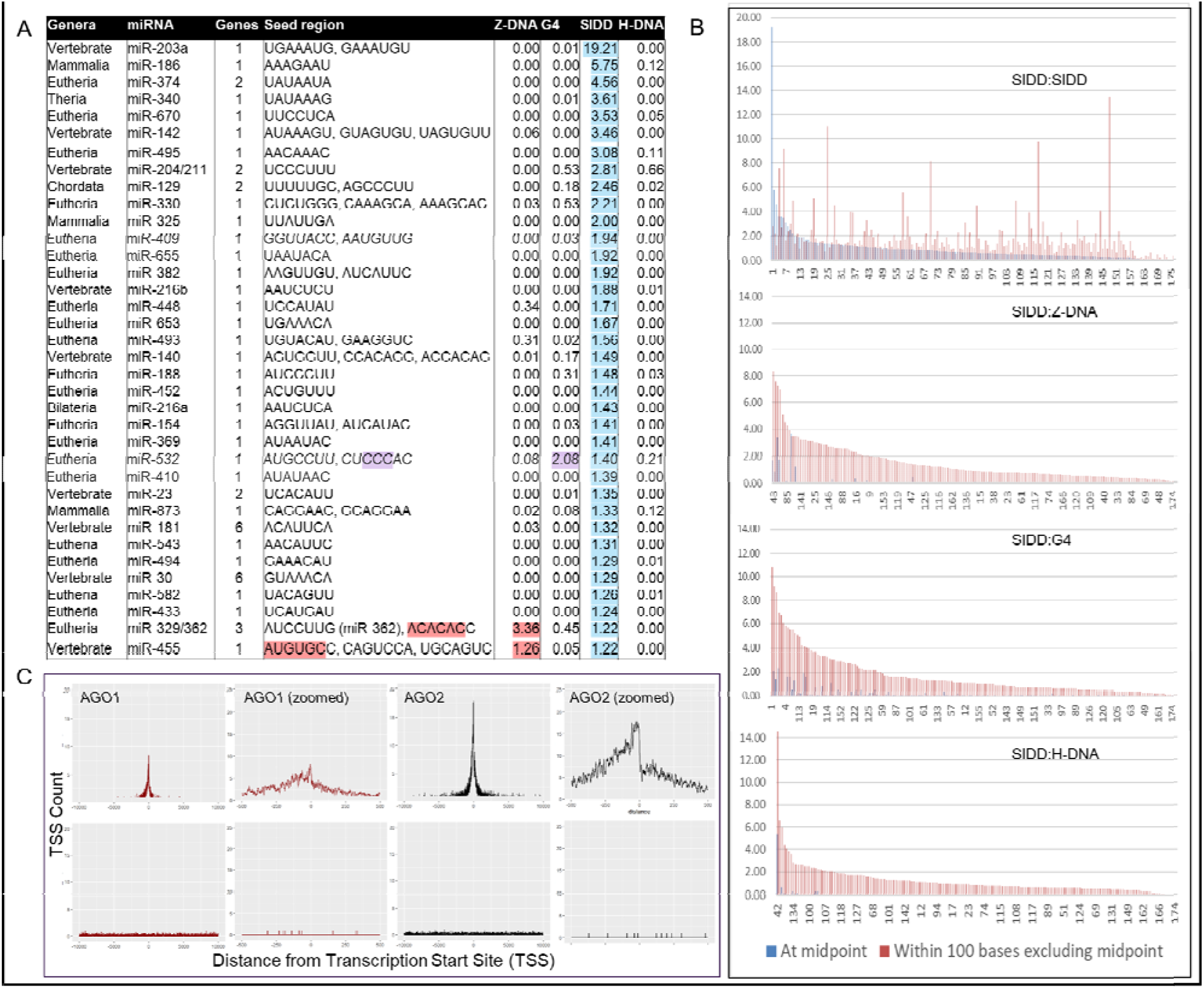
**A**. Mapping of highly conserved miRNA family seed sequences to SIDDs experimentally mapped in activated B-cells with the percentage of those that embed miRNA seed sequences highlighted in light blue. The SIDD sequences that overlap G- and Z-flipons are colored purple and primrose respectively **B**. Relative position of Z-DNA, G4 and H-DNA to SIDDs. The charts are ordered by frequency, not by miRNA family. The blue vertical bars show the percentage of SIDDs that have a complete overlap with the miRNA seed sequence. The red lines are the percentage of matches for experimentally determined Z-DNA, G4 or H-DNA within 100 bases either side of the SIDD/miRNA match. The additional SIDD matches in this region are for the same miRNA. C. The position of ENCODE AGO1 and AGO2 ChIP-seq peaks in HepG2 liver cells relative to the Transcription Start Site (TSS) is shown in the upper panel along with a zoomed in image for both. For comparison, the overlap with randomly selected sequences is shown in the lower panel. SIDDS located in promoters show a 15.05% and 25.47% overlap with AGO1 and AGO2 peaks respectively compared to 1.91% and 4.79% genomewide. The genomewide features recognized by each protein is shown in supplemental figure 1.

We tested the colocalization of experimentally determined SIDDs with other flipon classes by extending the search window to 100 bases either side of the initial SIDD/miRNA seed match (Figure 2B). We found some instances where the same miRNA seed sequence match was present at multiple sites within a region (top panel Figure 2B, the index interaction is shown with blue lines, red lines show additional matches). Allowing matches with other miRNA seeds yielded a potential miRNA binding site within 500 basepairs for 70% of flipons. We also found a high frequency of Z-, G- and T-flipons close to the index SIDD/miRNA seed match (Figure 2B, Supplementary Data 1).

The top hits for a Z-flipon match with a single miRNA seed sequences include miR-377, miR-744, miR-329/362 and miRNA-342 (8.34%, 7.63%, 7.25% and 7.00% overlap respectively), all containing alternating pyrimidine/purine Z-DNA prone sequences. These miRNAs potentially regulate Z-DNA formation by promoting R-loop formation. A similar situation occurs for G4-forming sequences with the top hits containing a run of three purines on one strand or the other, again suggesting a reciprocal relationship between flipon conformations. For experimentally validated H-DNA, the top hits include miR-185, miR-203a and miR-204/211, each with polypurine seeds and overlaps of 15.32%, 6.56% and 6.04% respectively. In most other cases, the match of the miRNA seed sequence to canonical Z-, G- and T-flipon sequences was low, suggesting that these elements are distinct from S-flipons. Indeed, the spectrum of colocalization of miRNA with differs by flipon class (the position of miRNAs named on the x-axis is quite variable) (Figure 2B).

To confirm the interaction of miRNAs with SIDDs, we examined the binding of AERs to DNA using the ENCODE AGO1 and AGO2 ChIP-seq datasets that are available for the human HepG2 liver cell line [35]. Their binding sites are highly enriched in promoters (70.17% and 54.68% overlap within 1 Kb of TSS respectively, Supplementary Figure 1) [35]. For SIDDs located in promoters, we found a 15.05% and 25.47% overlap with AGO1 and AGO2 peaks respectively compared to 1.91% and 4.79% for SIDDs located throughout the genome. The pattern of binding around TSS shows a wide distribution of AGO contacts consistent with multiple different miRNA binding sites (Figure 2C). By comparison, the overlap of AGO1 and AGO2 with the experimentally validated Z-flipons in promoters was 4.23% and 3.76% respectively, consistent with results from the observed overlap of seed and flipon sequences.

The close proximity between the different flipon classes was confirmed genomewide (Figure 3). The enrichment of experimentally determined Z- and G-flipons in promoters is consistent with previous results [14,20,21] (Figure 3A). Multiple exact matches for the same miRNA seed is frequent for SIDDs (41.01% Figure 3B and 67.93% when combined with single exact matches), suggesting that this feature is under selection. The tight association with SIDDs and other classes of flipons increases as the search window is expanded to 500 bp either side of the SIDD(Figure 3B). The pairing of Z-DNA with SIDDs in promoter regions is smaller numerically than with G4 and H-DNA but constitutes a higher percentage of total Z-flipon counts, increasing to 33% within 500 bps(Figure 3C and Supplementary Figure 2, 3). Gene ontogeny shows that the promoters identified in the 500 bp window are highly enriched for TF involved in development, particularly those involved in cell fate commitment (Table 1, Supplementary Data 2)

**Figure 3.**
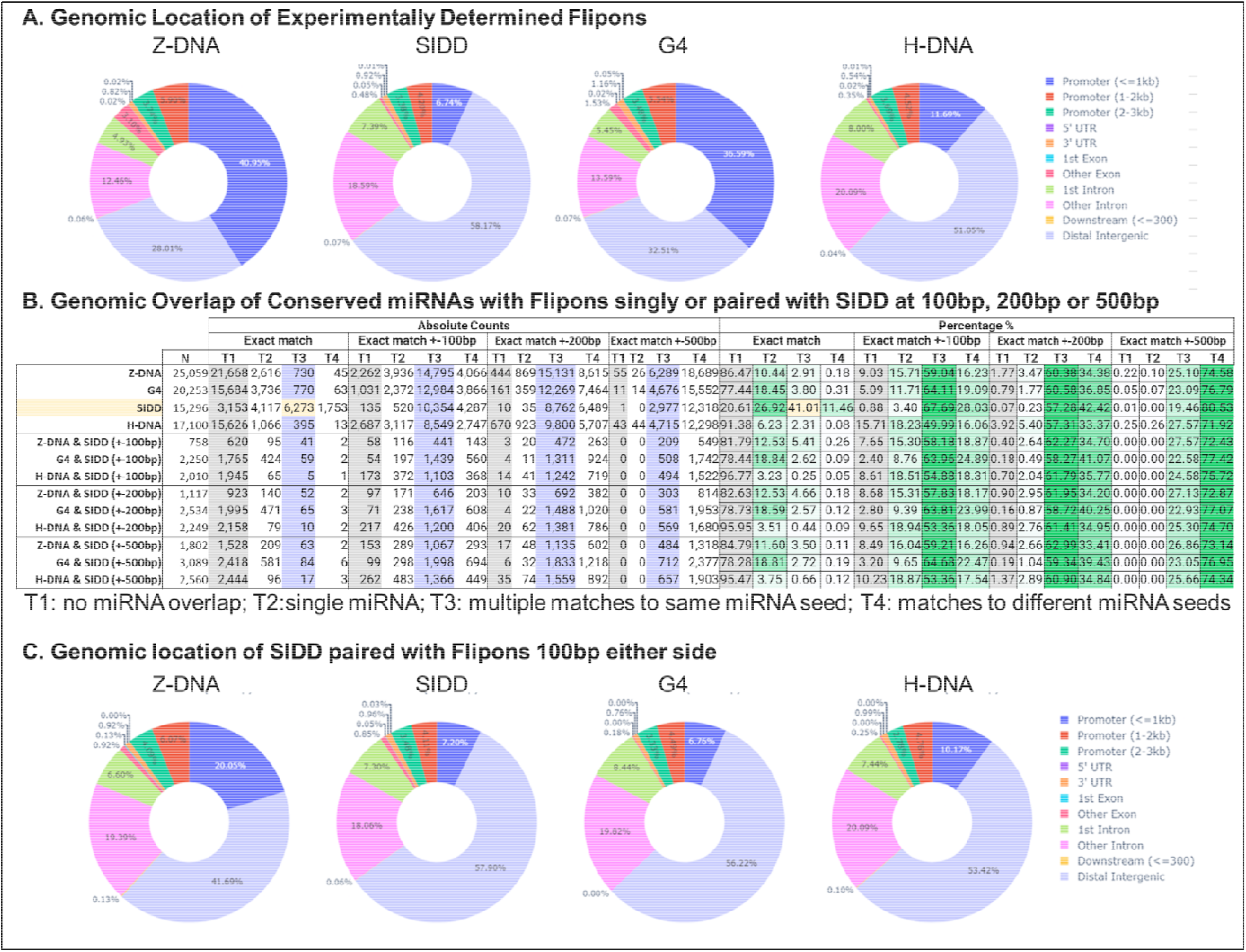
Mapping of Flipon overlaps to miRNAs and with SIDDs **A**. The genomewide distribution of experimentally mapped flipons in murine B-cells [22] **B**. The mapping of miRNA seed sequences to flipons either with an exact match or a match within 100, 200 or 500 basepairs of the initial match site. Grey indicates the number of miRNAs not mapped in the region scanned (T1). T2 represents a single match in the region. Lilac shows the counts for multiple overlaps in a region with the same miRNA seed (T3). Overlaps, such as those highlighted in cream, are frequent for SIDDs. T4 represents the number of overlaps with different miRNA seeds. Overlap percentages are highlighted in green. In addition, the colocalization of a flipon and SIDD within a region is counted along with and the miRNA seed sequence overlap. **C**. The genomic location of flipons within 100 basepairs of the closet SIDD is illustrated with respect to gene features.

## Discussion

The close proximity of SIDDs to other classes of flipons leads to the general model presented in figure 1 where the exchange of energy between the different conformations is modulated by the power stored within nucleosomes (Figure 1A and B). In Figure 1C, we show that the flip from Z-DNA to B-DNA during nucleosome docking transfers energy to the S-flipon, opening up the SIDD to allow sequence-specific binding of an RNA to form an R-loop. In panel 1D, we show a G-flipon storing energy that can be used by a helicase to place a nucleosome in a position to stabilize a triplex [28].

We found an enrichment of TF involved in cell fate specification in promoters where SIDDs were paired with other flipons (Table 1). To better understand how flipon pairs and miRNAs interact, we turned to early development where the sequence of events can be established by the time at which gene knockout is embryonically lethal. The time of death given in Table 2 is for mice homozygously deficient in key histone methyltransferases. The row ordering informs as to the epigenetic modifications made by each enzyme is necessary for a subsequent developmental step. The first step requires SETDB1 to suppress endogenous retroelements that are actively producing double-stranded RNAs [36,37] and results in the H3K9me3 dependent formation of constitutive heterochromatin [38]. The second critical step is the marking of CpG rich promoters with H3K4me3 by SETD1A [39]. The enzyme is guided by the CXXC1 protein and related family members that bind at CpG steps [40,41]. The lysine methylation likely facilitates the ejection of nucleosomes that potentially powers the flip of DNA to an alternative conformation. The third step is the creation BiPs, when H3K27me3 is written by the enhancer of zeste homolog 2 (EZH2) methyltransferase to sites marked by H3K4me3. EZH2 is part of the polycomb repressive complexes 2 (PRC2) and is essential for normal development [42]. The guidance of PRC2 by small RNAs is well established in worms and yeast but fewer examples are known in mammals [43-45]. The H3K27me3 is thought to protect against DNA methylation that accompanies heterochromatin formation[39]. The fourth step depends on protein factors that edit BiP marks to produce tissue-specific outcomes and involves PRC1 [46]. The diversity of PRC1 complexes suggest they have a number of divergent roles in tissue development [47]. However, deletion of the PRC1 enzyme EZH1 is not embryonically lethal [48].

**Table 2:** RNA-Directed DNA Transcription. Histone writers described in the text, listing, modification, lethality in homozygous null embryos and the chromatin feature they are associated with. (me: methyl; ub: ubiquitin; CBX: chromobox proteins; CXXC: Cysteines separated by two other amino acids; PRC: Polycomb Repressive Complex; SET: **S**u(var)3-9, **E**nhancer-of-Zeste and **T**rithorax; EZH: **E**nhancer-of-**Z**este; EHMT: Euchromatic histone-lysine N-methyltransferase; siRNA: small interfering RNA)

| Methyltransferase | Targeting | Modification | Null (Lethality) | Chromatin Feature | Outcome |
| --- | --- | --- | --- | --- | --- |
| SETDB1 | siRNA | H3K9me3 | E3.5-E5.5 [55] | Constitutive Chromatin | Suppression of retroelements [36,37] |
| SETD1 | CXXC1 | H3K4me3 | E6.5-E7.5 [40] | CpG Rich Regions Tag | Prevention of DNA methylation |
| PRC2(EZH2) | miRNA | H3K27me3 | E7.5-E10.5 [56] | Bivalent Promoters | Fate Specification |
| EHMT1, EHMT2 | Transcription Factors | H3K9me3 | E9.5-E12.5 [57] | Facultative Chromatin | Tissue Differentiation |
| PRC1(EZH1) | CBX proteins | H2AK119ub | Normal [48] | Active genes [58] | Cell-specific programs |

We posit that interaction of flipons with *trans*-RNA bootstraps the initial phase of embryogenesis by specifying sites destined to become bivalent promoters (Figure 4). The CpG rich regions in layer 1 are enriched for promoter flipons. The adoption of a non-B-DNA conformation is favored by H3K4me3 and ensures that TSS and SIDDs are nucleosome-free and in open chromatin. The *trans*-RNAs that bind SIDDs (layer 2) then specify the location of BiPs by promoting H3K27me3 (layer 3). These tags localize the cellular machinery needed for later steps involving sequence-specific binding by TF and editing of chromatin marks (layer 4). A subset of TF interactions may be further moderated by miRNAs that sequester their binding sites into R-loops or triplexes [49]. Each of the four layers utilizes different sets of information to genetically encode developmental programs. The initial specification of bivalent promoters is by the *trans*-RNAs transmitted to the embryo by each parent. These include those miRNA derived from ovary and spermatozoan specific miRNAs [50,51], transmissions with evident evolutionary ramifications [52].

**Figure 4.**
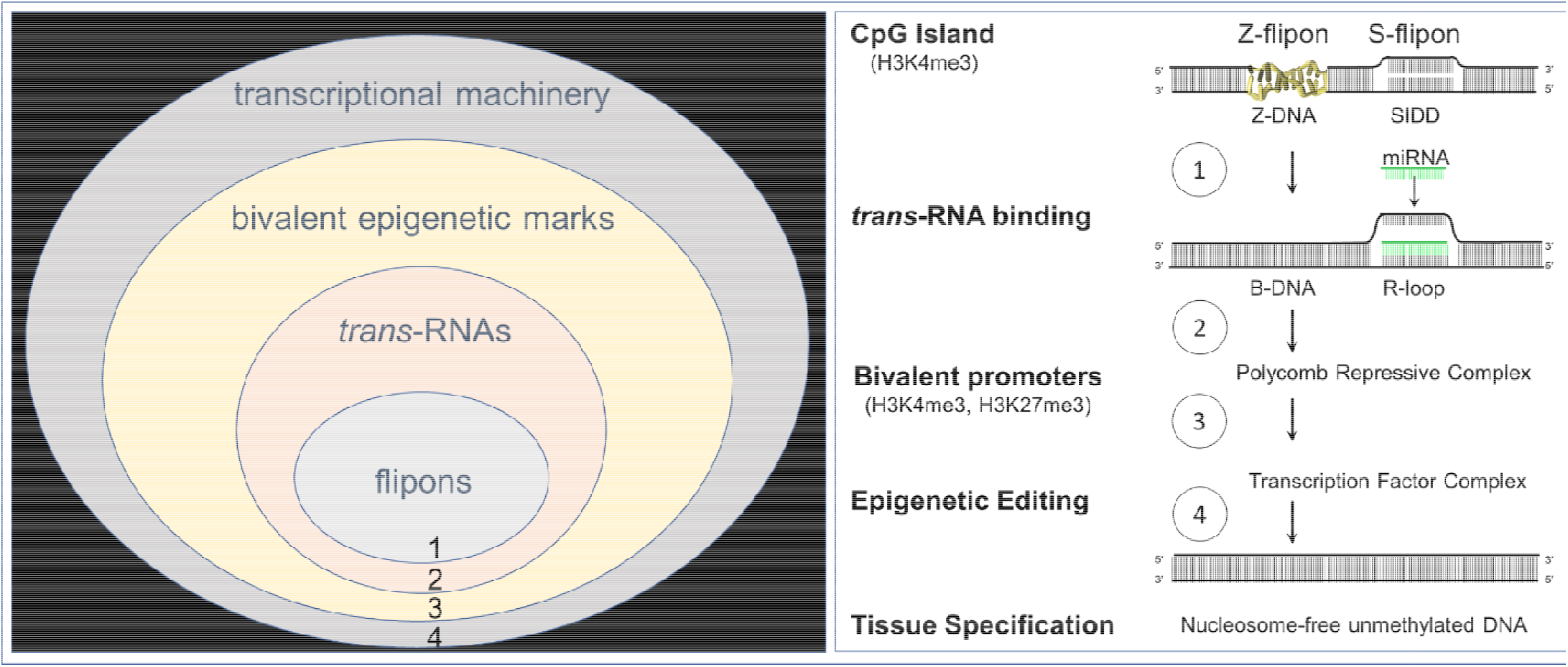
The four numbered layers proposed as necessary to bootstrap development of an embryo. Flipons and *trans*-RNAs have conjoint roles. Flipons, such as those that form Z-DNA and G4 quadruplexes, phase nucleosome to place transcription start sites (TSS) and *trans*-RNA binding sites in regions of open chromatin. Evolutionarily conserved miRNAs represent one class of *trans*-RNAs that recognize SIDDs (Stress-Induced Duplex Destabilization Sites) and form R-loops or triplexes in these AT-rich regions [26,27]. The SIDD sequences are referred to in the text as S-flipons. Early in development, we propose that the engagement of S-flipons by miRNA binding sites in close proximity to TSS establishes bivalent epigenetic tags that underlie the repressive and active states found later during tissue specification. The removal of one or other or both bivalent tags depends on the sequence-preferences of transcription factors and the enzymes present in the complex. Addition of further tags such as K2AK119ub further modify outcomes. Each of the four proposed regulatory layers coordinate development by accessing different types of genetically encoded information.

### Future Steps

The results we present are for a subset of conserved miRNAs from a single cell line and likely underestimates the true extent of these interactions that may also involve PIWI [8] and other *trans*-RNA types [2]. The SIDDs involved in specifying outcomes likely vary by tissue. Their detection by KMnO_4_ footprinting may be incomplete. This technique relies on thymine modification with other unpaired bases that can be detected with kethoxal missed [53]. Both footprinting techniques can be applied to any cell line and also to any early stage of embryonic development to provide more information on how flipons affect the readout of genetic information. The ability to control exposure and duration [22] permits a careful mapping of chronological events to establish cause and effect. The approach can also potentially inform on how miRNAs coordinate nuclear and cytoplasmic gene expression. Other potential applications include the targeting of lineage-specific S-flipons with RNAs that alter binding sites for both enhancer and repressor protein complexes [49].

## Methods

### Datasets

AGO1 ChIP-seq (GEO:GSE174905), AGO2 ChIP-seq (GEO: GSE136467), KMnO_4_ mapping of non-B-DNA structures (https://www.ncbi.nlm.nih.gov/CBBresearch/Przytycka/index.cgi#nonbdna), conserved miRNA seed sequences(Bartel [4])

### Code Availability

https://github.com/TheodorrodeohT/article_conserved_miRNA.

The following types of flipons were uplifted from mm9 to mm10 using UCSC liftOver. ‘bedtools 2.26’ software was used to find overlaps between ‘.bed’ files and apply slops of different sizes to the initial flipon coordinates. The input ‘.bed’ files were converted into ‘fasta’ format using ‘biopython 1.79’ python library. Each flipon sequence was tested for embedding miRNA seed sequences via default python string and array operations. The results were saved as a ‘pandas’ dataframe and later converted into ‘.xlsx’ format for better readability. Values for pie charts were calculated using ‘ChIPpeakAnno’ and ‘ChIPseeker’ R libraries. The genomic annotation used as a reference was [GENCODE vM25] (https://www.gencodegenes.org/mouse/release_M25.html) (GRCm38.p6). The pie charts were drawn using ‘plotly 5.8.0’ python library.”

For the analysis of AGO1 and AGO2, we used custom R (4.1.3) and python (3.7.13) scripts. The scripts are available in the repository https://github.com/Madund3ad/article-mi-RNA/. The genomic annotation used as a reference was [ENSEMBL GRCh37.87 http://anonymous@ftp.ensembl.org/pub/grch37/release-87/gtf/homo_sapiens/Homo_sapiens.GRCh37.87.gtf.gz (GRCh37.87) Overlaps and distances to TSS were calculated using” GenomicRanges” R library and values for smooth curves were calculated using the “zoo” R library. After that distances to TSS were drawn using “ggplot2” R library. Pie charts’ data were calculated using ‘ChIPpeakAnno’ and ‘ChIPseeker’ R libraries and drawn using ‘plotly 5.8.0’ python library.

## Analysis

The ssDNA sequence motifs of non-B DNA from Kouzine paper are used as the peak data set. Since the original version of the mouse genome was mm9, the peaks were uplifted to mm10 using the UCSC liftOver utility (http://hgdownload.soe.ucsc.edu/admin/exe/linux.x86_64/), resulting in slightly different N dimensions of each dataset: H-DNA (17,110 → 17,000), G4 (20,260 → 20,253), Z-DNA (25,063 → 25,059), SIDD (15,299 → 15,296). Using Python 3.7 and biopython 1.79 library, the peak coordinates were converted into nucleotide sequences. Then, each sequence was tested for overlap with each miRNA seed region. The final results are compiled into a summary table for comparative analysis of the overlap values. Gene enrichment was performed using the WEB-based GEne SeT AnaLysis Toolkit (http://www.webgestalt.org/).

For the analysis of distance to TSS distribution we used ENCODE annotation (GRCh37.87). Only the distance to the nearest region of interest (AGO1 or AGO2 ChIP peak) is shown on the plot. A rolling average with a window of 8 bases is applied to smooth the curve.

For the analysis of overlaps between ENCODE AGO narrow peaks, experimental Kouzine SIDD peaks and promoters we took 1000bp upstream regions from TSS as a promoter. TSS coordinates were obtained from ENCODE annotation (GRCh37.87). Two regions were considered to be overlapping if they had at least one base pair in common. Significance of the overlap was calculated using Monte-Carlo simulation (n=1000).

## Contributions

AH conceived and wrote the paper with input from MP, FP and DK. FP analyzed the interactions between flipons and miRNAs and DK analyzed the locations of AGO1 and AGO2 in K562. MP helped edit the paper and supervised the analysis by FP and DK.

## Acknowledgements

We thank Sid Balachandran for reviewing the manuscript

## Conflicts of Interest

AH is the founder of InsideOutBio, a company that works in the field of immuno-oncology. All the authors declare that the research was conducted in the absence of any commercial or financial relationships that could be construed as a potential conflict of interest.

**Supplementary Figure 1.**
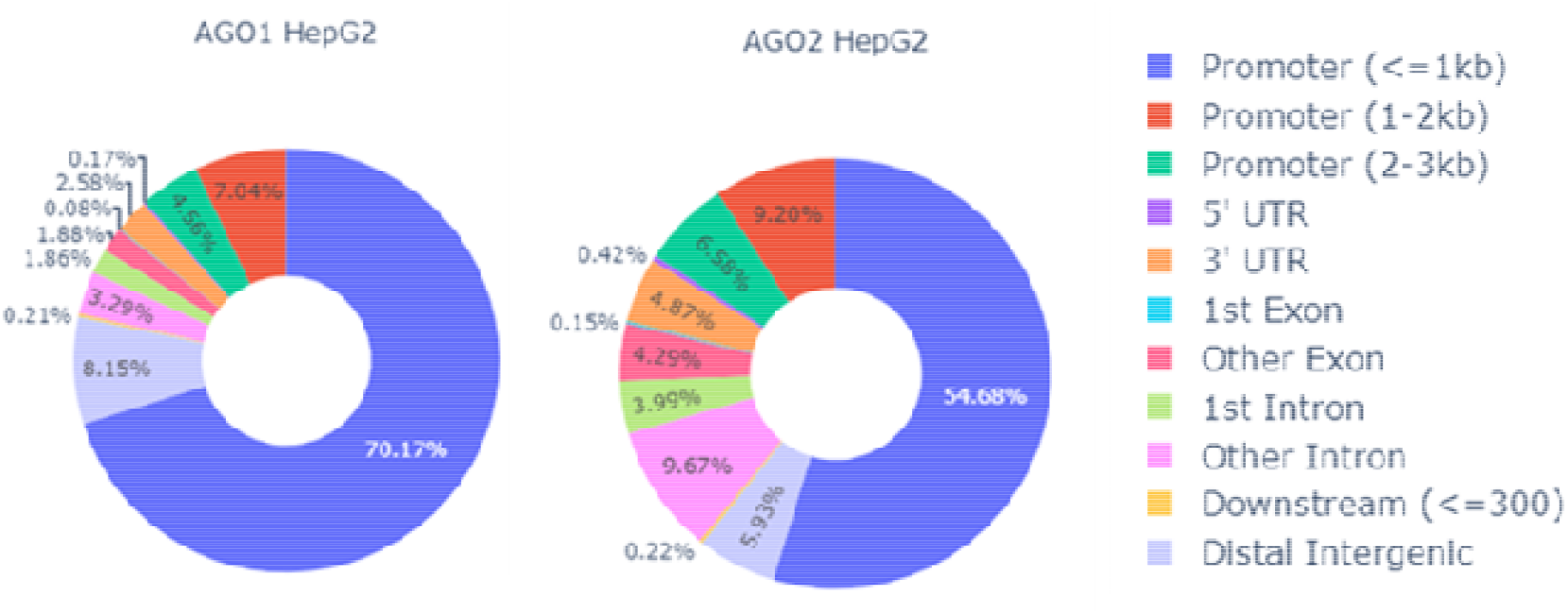
The genomic location of ChIP-seq peaks for AGO1 and AGO2 in human HepG2 liver cells annotated by feature.

**Supplementary Figure 2.**
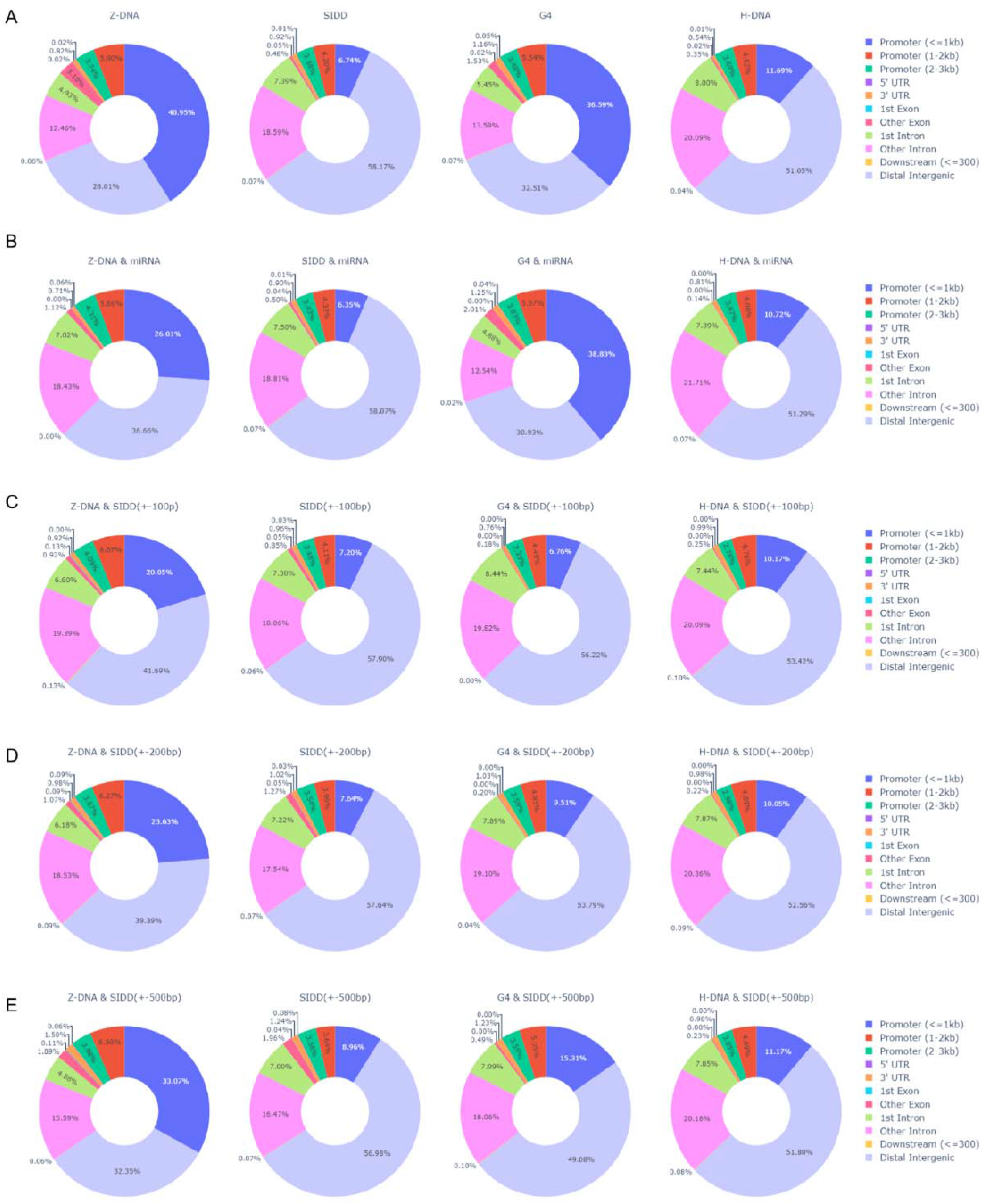
The location of flipons within genomic features. **A.** Mouse genome **B**. Location of flipons matching a conserved miRNA seed **C**. Flipons within 100 basepairs of a SIDD **D**. Flipons within 200 basepairs of a SIDD **E**. Flipons within 500 basepairs of a SIDD.

**Supplementary Figure 3.**
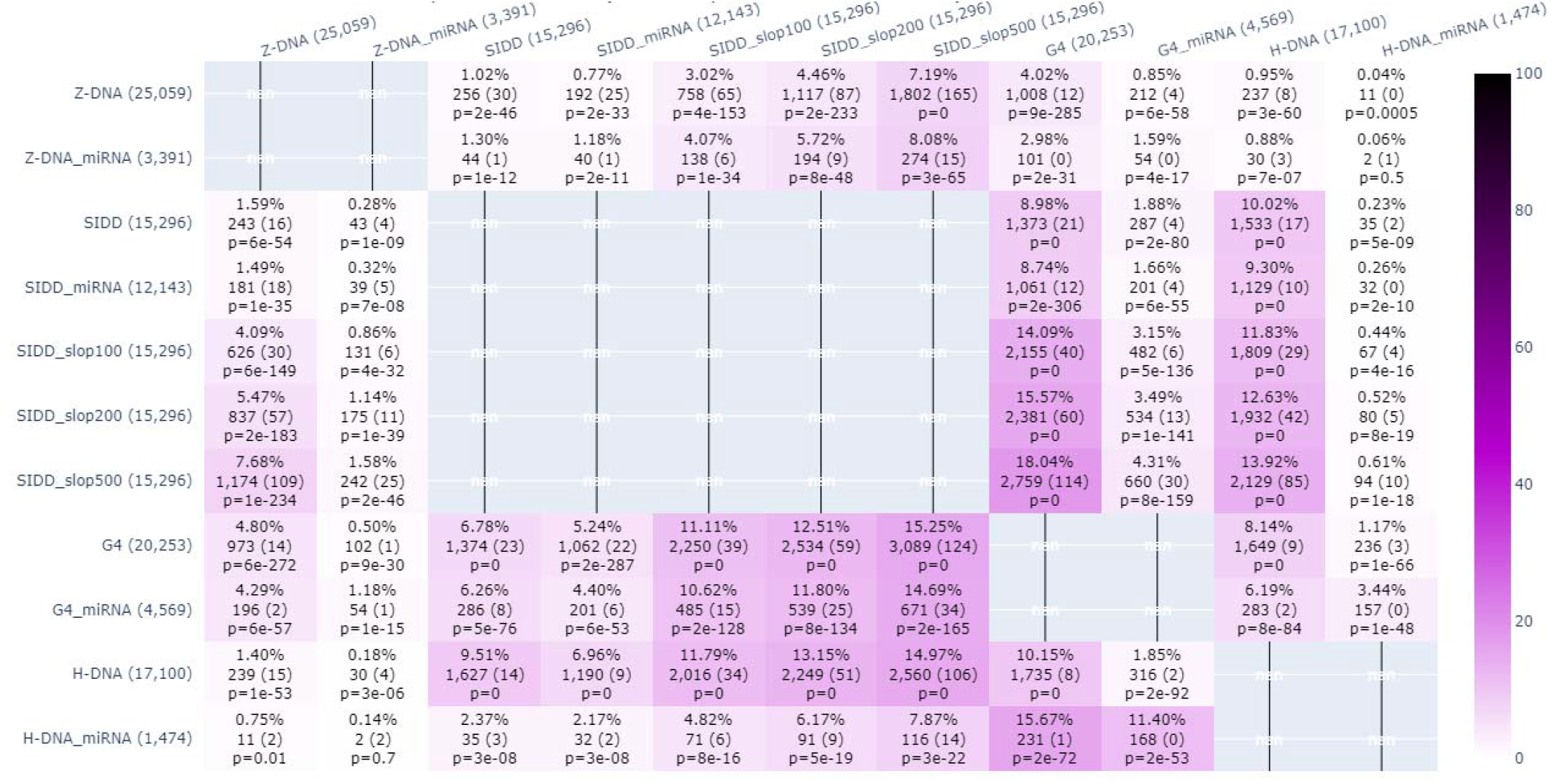
Enumeration of the overlap at different distances between the different features examined in the KMnO4 footprinting dataset [22].

